# *In vitro* model of inflammatory, hypoxia, and cancer stem cell signaling in pancreatic cancer using heterocellular 3-dimensional spheroids

**DOI:** 10.1101/454397

**Authors:** Megha Suresh, George Mattheolabakis, Amit Singh, Mansoor Amiji

## Abstract

**Introduction:** As one of the most aggressive cancers worldwide, pancreatic cancer is associated with an extremely poor prognosis. The pancreatic tumor microenvironment consists of cancer cells and other tumor associated cells. Cross-talk between these different cell types through various signaling molecules results in the development of a more aggressive and malignant phenotype. Additionally, due to the highly dysregulated vasculature of tumors, the inner tumor core becomes hypoxic and eventually necrotic. Therefore, there is a need for the development of a physiologically relevant *in vitro* model that recapitulates these dynamic cell-cell interactions and the 3-dimensional (3D) structure of pancreatic tumors.

**Methods:** Four different 3D co-culture spheroid models using different combinations of Panc-1 tumor cells, J774.A1 macrophages, and NIH-3T3 fibroblast cell lines were reproducibly developed using the hanging drop technique in order to mimic the tumor microenvironment and to evaluate the differences in expression of various inflammatory, hypoxia, and cancer stem cell markers, including IL-8, TNF-α, TGF-β, HIF-1α HIF-2α, SCF, and LDH-A. Additionally, immunofluorescence studies were employed to investigate whether these spheroids tested positive for a cancer stem cell population.

**Results:** Pronounced differences in morphology as well as expression of signalling markers were observed using qPCR, indicative of strong influences of co-culturing different cell lines. These models also tested positive for cancer stem cell (CSCs) markers based on immunofluorescence and qPCR analysis.

**Conclusion:** Our results demonstrate the potential of 3D co-culture spheroid models to capture the inflammatory and hypoxic markers of pancreatic tumor microenvironment. We further demonstrate the presence of cancer cells with stem cell markers, similar to actual pancreatic cancer tumor. These spheroids present excellent *in vitro* system to study tumor-immune-stromal cell interactions as well as test deliverability of potential therapeutics in the tumor microenvironment with accurate physical and physiological barriers.

## 1. Introduction

Pancreatic cancer is one of the deadliest cancers associated with the 4^th^ highest number of deaths amongst all cancers worldwide, with the incidence rate nearly equaling the mortality rate. In 2018, it is estimated that 55,440 adults will be diagnosed with pancreatic cancer and 44,330 patients will die of the disease (1). With the growing epidemic of type I diabetes worldwide, the number of patients with pancreatic cancer is expected to increase significantly in the coming years. Despite tremendous efforts to improve patient survival, including surgical resection, chemotherapy, and radiation therapy, these options have not been successful in prolonging patient life beyond a few months in pancreatic cancer (1-5). The poor prognosis in pancreatic cancer patient is largely associated with late presentation of the symptoms when the disease has already progressed to advanced stage where significant metastasis has occurred combined with lack of effective therapies that can improve patient outcomes. Additionally, there is a need to develop suitable *in vitro* and *in vivo* disease models that can improve the drug development process.

The pancreatic tumor microenvironment is represented by an outer proliferative and inner necrotic zone, along with stromal components including the extra-cellular matrix, blood vessels, signaling molecules, and other cells including tumor-associated macrophages (TAMs), tumor-associated fibroblasts (TAFs), and endothelial cells form (6, 7). Animal models for pancreatic cancer recapitulate the tumor microenvironment to some extent but are difficult to develop, time consuming and very expensive (8). Two dimensional (2D) cell monolayers are simple to culture and provide convenient testing platforms for screening anti-cancer drugs but they are not truly representative of the tumor microenvironment, morphologically or functionally (9). More recent efforts have shifted focus on three dimensional (3D) co-culture spheroids that serve as a robust in vitro model that exhibit several features of pancreatic cancer microenvironment (10).

Within a 3D environment, cancer cells can switch to a stem cell like phenotype, which is more aggressive, metastatic and shows higher resistance to chemotherapy. Additionally, these cancer stem cells or CSCs are pluripotent, repair damaged cells and show self-renewal characteristics. In other words, a CSC can divide to form one daughter CSC and one non-tumor initiating cell. However, it is unclear where these CSCs originate from, if tumor cells develop into CSCs or if they are formed from stem cells. These cells are characterized by various markers, such as stem cell factor (SCF), epithelial surface antigen (ESA), CD24, CD44 and CD133 (11-13).

Hypoxia regulates tumor cell phenotype and cancer cells adapt their metabolism to sustain their growth, unlike healthy cells that are severely damaged in hypoxic conditions. Cancer cells can also undergo genetic mutations under hypoxia to give rise to multi-drug resistance (MDR), enhanced angiogenesis and migration. Cancer stem cells regulate their microenvironments to form niches, within which they reside. CSCs, in fact, prefer hypoxic environments because they avoid DNA damage from reactive oxygen species at higher oxygen levels. Under low oxygen conditions, levels of hypoxia inducible factors, HIF1α and HIF2α are elevated. However, CSCs tend to up-regulate HIF2α levels regardless of oxygen levels, to induce an artificial hypoxic environment within which they thrive, sustain their self-renewal properties and maintain their undifferentiated state (14-19).

The dynamic interplay among cancer, stromal and immune cells occurs through various cytokines and chemokines. Numerous signaling pathways and transcription factors, including NF-κB, Snail and Slug, drive CSC formation as well as epithelial-mesenchymal transition (EMT). IL-6, TNF-α and IL-1β bring about HIF-1α release, activating the NF-κB pathway, which ultimately leads to the down-regulation of apoptotic genes. Snail and Slug are also activated, regulating E-cadherin levels, thereby driving EMT (20-22). Sonic hedgehog (Shh) pathway activation, induced by TNF-α, leads to increased tumor cell proliferation through Snail and other EMT regulators (23, 24). Activation of the tumor suppressor gene p53 causes high mobility group box 1 (HMGB1) release, leading to TNF-α release from neighboring immune cells, inducing NF-κB and Snail activation (25, 26). This leads to activation of CXCL1, which attracts myeloid cells towards the tumor, further facilitating EMT (27). Inhibition of NF-κB decreases EMT and proliferation rates of CSCs (23).

Transforming growth factor-beta (TGF-β), an EMT regulator, recruits immune cells to the tumor microenvironment, which then release pro-inflammatory cytokines, including tumor necrosis factor-alpha (TNF-α). With TNF-α also increasing TGF-β levels, a positive feedback loop is created (26, 28, 29). However, the CSCs themselves efficiently regulate the inflammatory state by releasing interleukin-8 (IL-8), MCP-1 and RANTES and bring about stromal cell proliferation and inflammation (30).

We have developed a single cell (homocellular) and 3-in-1 heterocellular spheroid model formed by self-association of Panc-1 human pancreatic adenocarcinoma cells, J774.A1 murine macrophages and NIH/3T3 murine fibroblasts that better recapitulate the 3D component as well as the cell-to-cell cross-talk in the microenvironment of avascularized tumors in early stages of growth. They also mimic the hypoxic gradient from the periphery to the core of the tumor (31). Therefore, they are crucial in helping gain a deeper insight into tumor biology as well as serve as an efficacious platform for testing of anti-neoplastic agents, thereby providing higher odds of success in *in vivo* testing of agents (32).

## 2. Materials and Methods

### 2.1. Cell lines and culture conditions

The cell lines used in this study include the human pancreatic epithelioid carcinoma - Panc-1 cell line, the J774-A1 murine macrophage cell line and the NIH/3T3 murine fibroblast cell line. All cell lines were obtained from American Type Culture Collection (ATCC, Manassas, VA). Panc-1 and J774.A1 cells were grown in 1X Dulbecco’s Modified Eagle’s Medium (DMEM) with 4.5g glucose and L-glutamine without sodium pyruvate obtained from Corning Cellgro (Manassas, VA). The growth medium was supplemented with 10% fetal bovine serum and 1% penicillin-streptomycin. NIH/3T3 cell line was grown in 1X DMEM supplemented with 10% fetal calf serum and 1% penicillin-streptomycin. All cell lines were cultured at 37°C at 5% CO_2_.

### 2.2. Establishment of homocellular and heterocellular 3D tumor spheroids

For the spheroid development, 96-well hanging drop plates from 3D-Biomatrix were used to grow single cell (i.e., homocellular) and multiple cell (i.e., heterocellular) spheroids. The spheroids were generated using 40μl of cell suspensions containing 20,000 cells in each well. For the homocellular spheroids, 20,000 Panc-1 cells were seeded in each well. For the Panc-1: J774.A1 and for the Panc-1: NIH/3T3 heterocellular spheroids, a 40 µl mixture of 10,000 tumor cells and 10,000 immune/stromal cells were seeded in each well. For the Panc-1: J774.A1: NIH/3T3 spheroids, a cell suspension mixture containing a 1:1:1 ratio of cell numbers with a total of 20,000 cells per well were seeded. We cultured spheroids for 5 days post-seeding and harvested them for downstream analyses. In order to maintain humidity in the spheroid plates, sterile 1X PBS was added to the in-built reservoirs on the 3^rd^ day of growth.

To measure the size of these spheroids, they were washed with 1X PBS upon harvesting, fixed with 4% Formalin and mounted on microscope depression slides (Fisher Scientific, PA) using the Shandon Immu-Mount (Thermo Scientific, PA), and covered with a cover-slip. The Carl Zeiss LSM 700 confocal microscope was used to capture images of spheroids and the Image J software was then used to measure and record the diameter. 5μm optical slices of spheroids were imaged through Z stacking on the confocal microscope and the total number of slices per spheroid was determined. This information was then used to calculate and record spheroid depth.

### 2.3 Distribution of Panc-1, J774.A1 and NIH3T3 cells within the heterocellular spheroids

In order to assess the arrangement of different cell types within the 1:1:1 heterocellular spheroid, Panc-1, J774.A1 and NIH3T3 cells were stained with Cellbrite Blue Cytoplasmic Membrane Labeling Kit, Cellbrite Green and Cellbrite Red Cytoplasmic Membrane Labeling Dye (Biotium, CA), respectively. 1X PBS was used to wash off the excessive dye, and the cells were re-suspended in 1X DMEM, counted and seeded to form spheroids using the method described in 2.2.

The spheroids were harvested on day 5 of growth, washed with 1X PBS, fixed with 4% formalin and mounted on a microscope depression slide using Shandon Immu-Mount. The slides were protected with a cover slip and fluorescent images were captured using the confocal microscope.

### 2.4. qPCR analysis of inflammatory, hypoxia and cancer stem cell signaling in spheroids

To assess the levels of various inflammatory, hypoxia and cancer stem cell markers, total RNA from homocellular and heterocellular spheroids was extracted using the Quick-RNA™ MiniPrep RNA extraction kit (Zymo Research, CA). The NanoDrop (Thermo Scientific, Willington, DE) was used to measure and record RNA concentrations. 1μg of total RNA were used for cDNA synthesis using the Verso cDNA synthesis Kit (Thermo Scientific, PA).

RT-PCR was used for the evaluation of inflammatory markers: IL-8, TNF-α, TGF-β. The Platinum PCR SuperMix (Life Technologies, NY) was used and both human and mouse primers, as shown in table 1, were used in order to ensure specie specific amplification with β-actin as the housekeeping control. The PCR products were run on a 2% Agarose E-Gel with SYBR Safe (Life Technologies, NY). The ChemiDocTM XRS imaging system (Bio-Rad, Hercules, CA) was used to image the bands. The intensity of the bands obtained for human IL-8, TNF-α and TGF-β were expressed as a percentage of the human β-actin band intensity for the respective samples. Similarly, the intensity of the bands obtained for the murine genes of IL-8, TNF-α and TGF-β were expressed as a percentage of the murine β-actin band intensity for each sample.

**Table 1.**
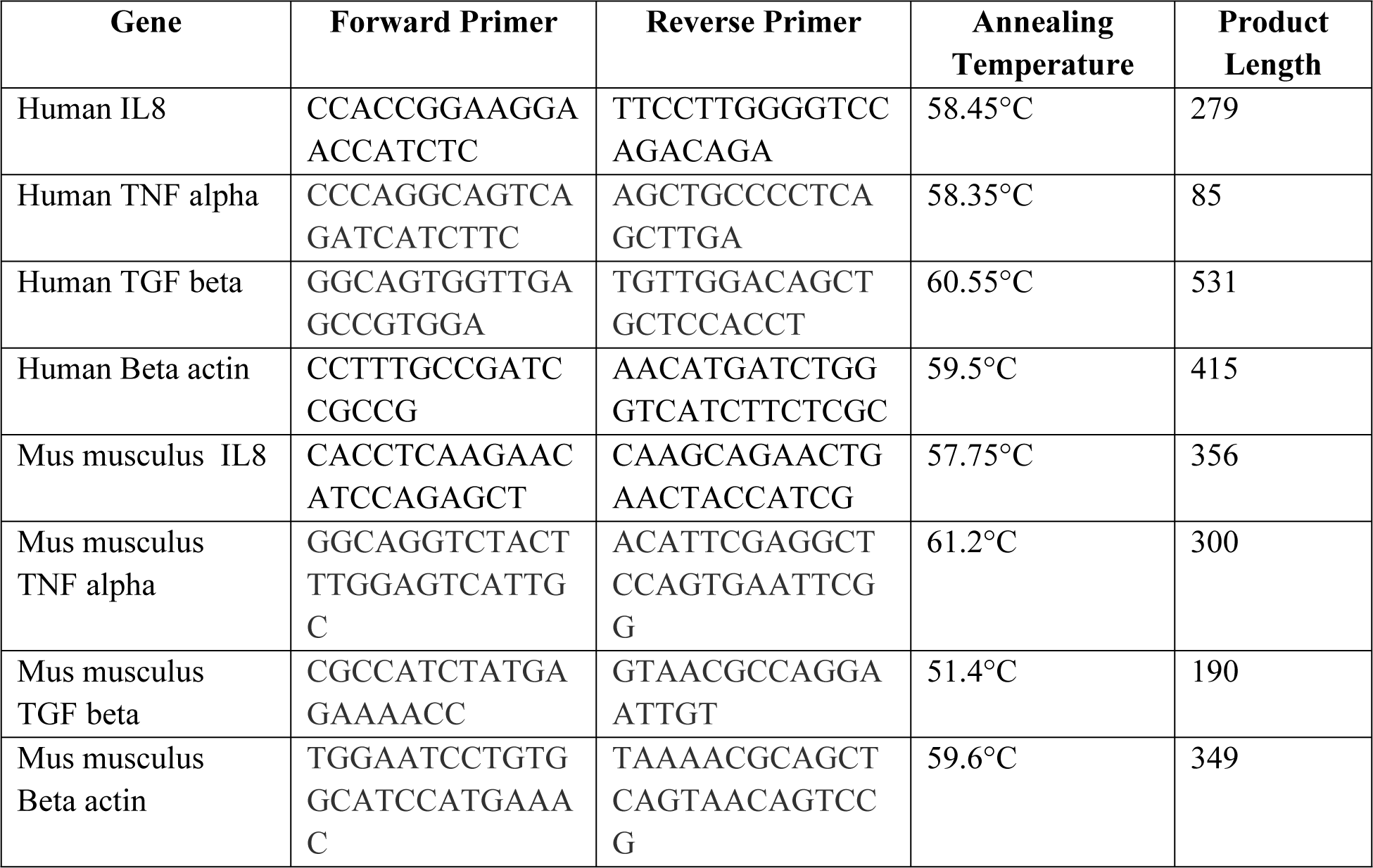
Human and Mouse Primer sequences for IL8, TNF alpha, TGF beta and Beta actin.

To evaluate levels of hypoxia markers: LDH-A, HIF-1α, HIF-2α, and cancer stem cell marker: SCF in normoxic cells, hypoxic cells, homocellular and heterocellular spheroids, the LightCycler^®^ 480 SYBR Green I Master kit (Roche Diagnostics, Indianapolis, IN) was used to carry out qPCR along with the Roche Light Cycler 480 machine. The primers used are listed in Table 2. Beta-actin was used as the housekeeping control, ΔCt values were obtained by calculating the difference between the Ct values of the target gene and beta actin of each sample and the 2 ^−ΔCT^ method of analysis was used to determine % gene expression.

**Table 2.**
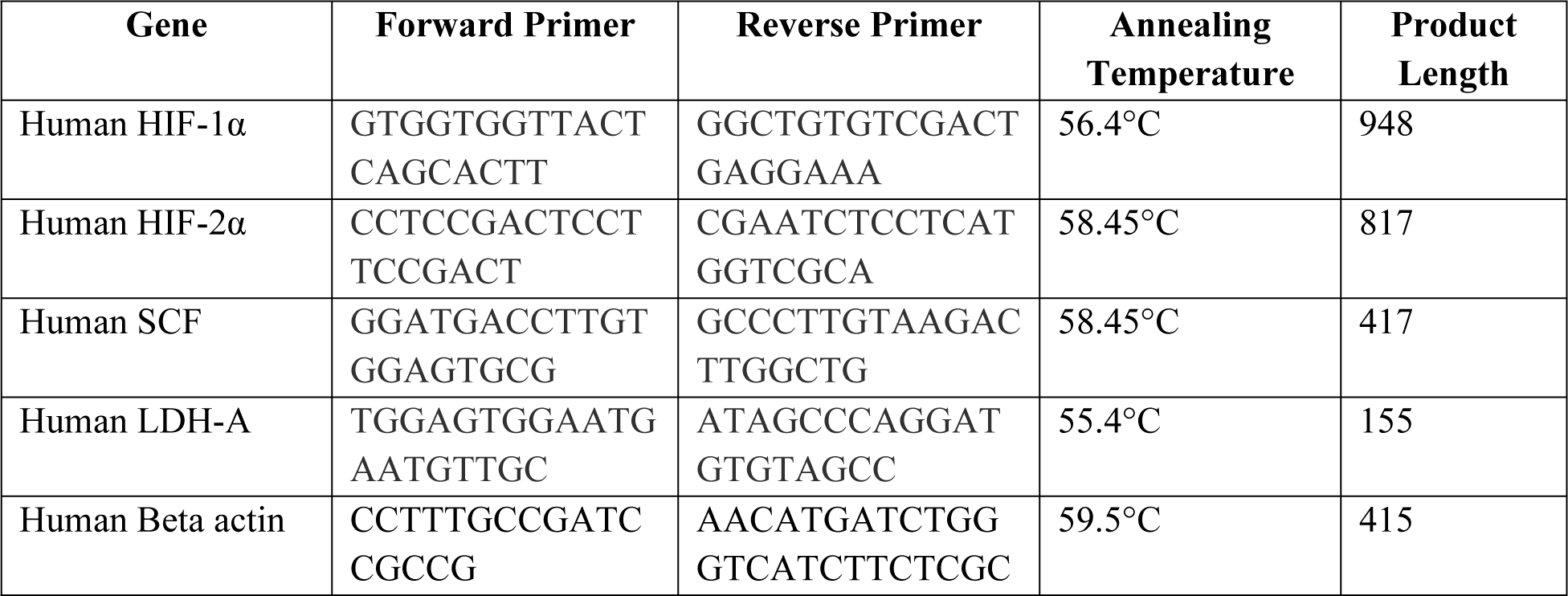
Human primer sequences for human HIF-1α, HIF-1β, SCF, LDH-A and β-actin.

### 2.5. Analysis of CD24+ and stem cell factor expression in spheroids

Spheroids harvested from hanging drop plates 5 days post-seeding were transferred to a 96-well Polystyrene assay plate (Corning Incorporated, NY) and 0.2ml Cell Staining buffer (BioLegend, CA) was added to each well to prevent receptor internalization. This was followed by fixation with 4% formalin. Spheroids were then washed twice with ice cold PBS, and blocking was carried out using a 10% FBS in PBS solution at room temperature for 30 minutes.

Spheroids were incubated for 10 hours with FITC mouse anti-human CD24 (BD Pharmingen, CA, catalog no. 55524, clone ML5) antibody or the rabbit polyclonal anti-human SCF primary antibody (Thermo Scientific, PA, catalog no. PA511563) in 2% BSA (bovine serum albumin, Fisher, PA) in 1X PBS. The SCF-stained spheroids were then incubated with a goat Anti-Rabbit IgG DyLite 594 conjugated highly cross adsorbed antibody diluted in 2% BSA in 1X PBS for 2 hours. All spheroids were then washed, and incubated with Hoechst 33342 Dye (Life Technologies, PA) to stain the nucleus, washed and imaged using the confocal microscope.

## 3. Results and Discussion

Spheroids were successfully grown using the 96-well hanging drop plates. The Panc-1 homocellular spheroids were the largest with a mean diameter of 919µm and an average thickness of 135µm, with a spherical morphology. The Panc-1:J774.A1 heterocellular spheroids were loosely formed spheroids with a mean diameter of 745 µm and an average thickness of 60µm and were also observed to be non-uniform in shape. The Panc-1:NIH/3T3 heterocellular spheroids were irregularly shaped with a mean diameter of 919.51µm and an average thickness of 116 µm, and the 3-in-1 heterocellular spheroids with a mean diameter of 396 µm and an average thickness of 55 µm formed the smallest and densest spheroids. The size and morphology of all four spheroid models is shown in Figure 1.

**Figure 1:**
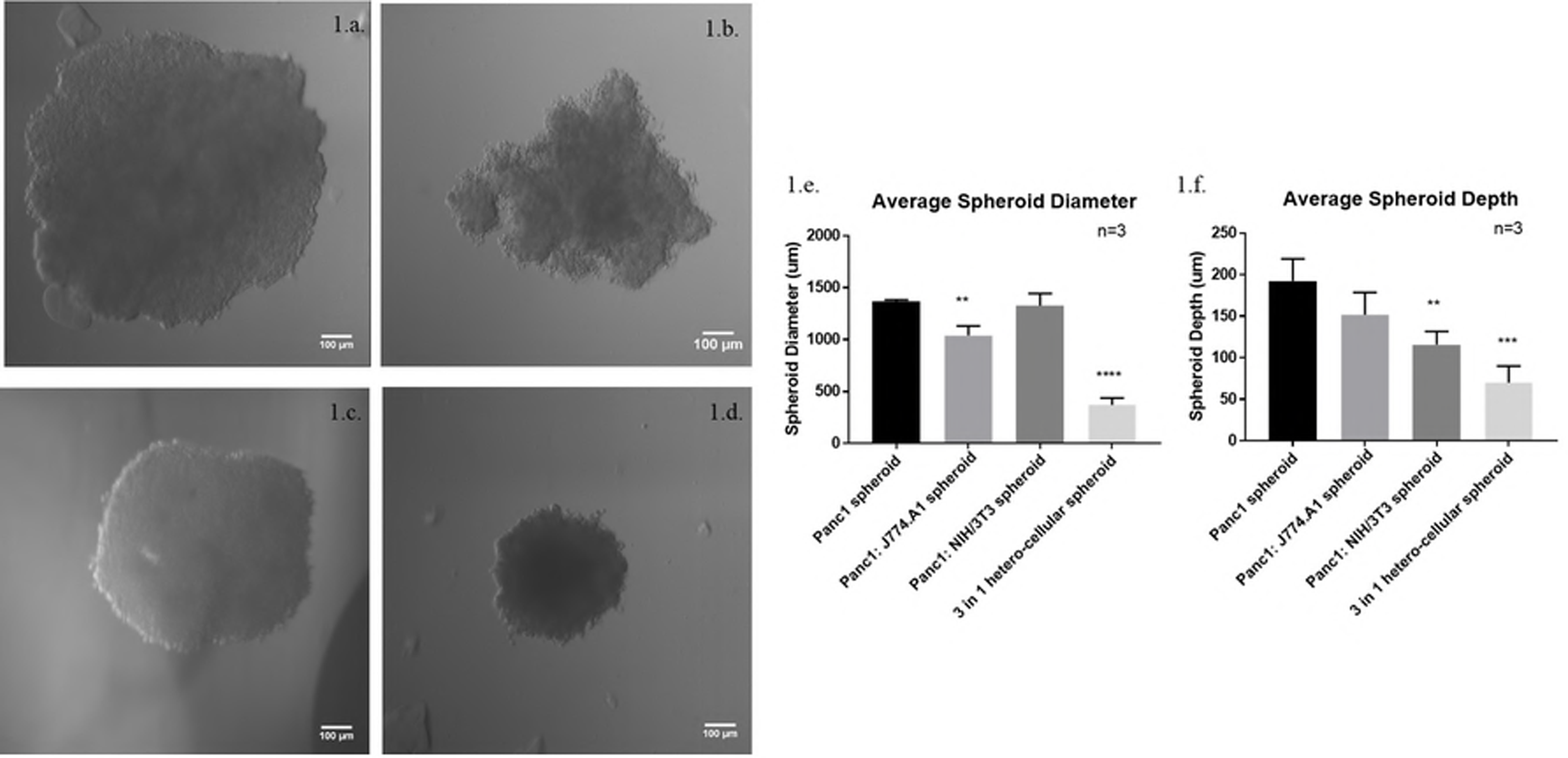
Morphological assessment of spheroids. A. Day 5 Panc-1 homocellular spheroid under 10X magnification - Average Diameter = 1365 microns and Average Thickness = 193 microns; B. Day 5 1:1 Panc-1:J774.A1 heterocellular spheroid under 10X magnification - Average Diameter = 1039 microns and Thickness = 152 microns; C. Day 5 1:1 Panc-1: NIH/3T3 heterocellular spheroid under 10X magnification - Average Diameter = 1327 microns and Thickness = 116 microns; D. Day 5 1:1:1 Panc-1: J774.A1: NIH3T3 heterocellular spheroid under 10X magnification - Average Diameter 371 microns and Average Thickness = 70 microns; E. Graphical analysis of spheroid diameter (n=3); F. Graphical analysis of the spheroid depth or thickness.

We stained the three individual cell types in 3-in-1 heterocellular spheroids with specific stains to understand their natural arrangement within the spheroids. As shown in Figure 2, Panc-1 cells were stained blue, J774.A1 cells in green and NIH/3T3 cells were stained in red. As evident from the images, no specific arrangement of the three cell lines was observed within the 3-in-1 heterocellular spheroid. This indicates that all cell types are in proximity of each other, which facilitates increased susceptibility to cross-talk through cytokine signaling. Giant cells were observed at the periphery of the spheroid. Since these were stained green, we hypothesize that these were formed by fusion of the macrophages. Molberg *et. al* carried out histological and immune-histochemical studies on ten pancreatic neoplasms and observed osteoclast-like giant cells in seven of these tumors. Their studies led them to conclude that the giant cells were histiocytic and non-neoplastic in origin (33). Multi-nucleated foreign body giant cells formed as a result of fusion of macrophages have been noted in thyroid, breast, skin and hepatic carcinomas as well, presumably drawn to the tumors in response to chemotactic factors and cytokines (34-37). However, immunocytochemical experiments employing epithelial, histiocytic and endothelial markers are required to confirm this hypothesis.

**Figure 2:**
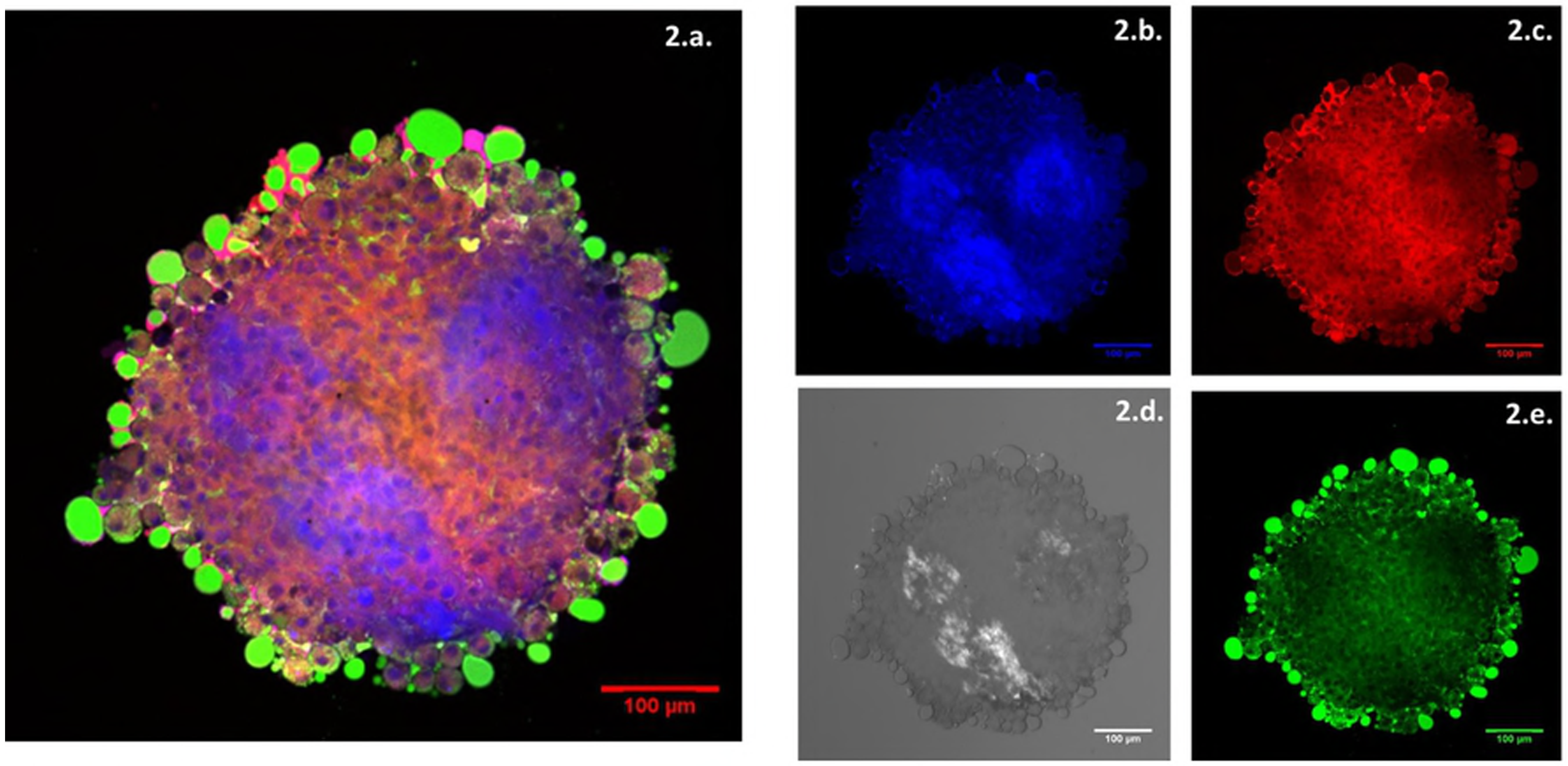
Arrangement of different cell lines within 3-in-1 heterocellular spheroids. A. Composite image showing distribution of Panc-1, J774.A1, NIH3T3 cells for the Day 5 1:1:1 co-culture spheroid; B. Panc-1 cells stained with Cellbrite Blue; C. NIH/3T3 fibroblasts stained with Cellbrite Red; D. Brightfield image of heterocellular spheroid; E. J774.A1 macrophages stained with Cellbrite green.

Next, we studied the expression of inflammatory, hypoxia and stem cell markers across different monolayer and spheroid models in order the assess the contribution of and interaction between different cell types in modulating the microenvironment. Analysis of inflammatory marker expression through RT-PCR and gel electrophoresis revealed distinct expression profiles across the spheroid models themselves and when compared to cells grown in normoxic and hypoxic conditions. Marker levels were expressed as a percentage of beta actin in the respective samples. IL-8 has been known to drive tumor metastasis, proliferation and survival and tumor cells undergoing EMT secrete increased levels of the pro-inflammatory cytokine (38). Additionally, senescent TAFs known to be up-regulated in pancreatic tumor microenvironments and TAMs secrete IL8 (39, 40). Highest levels of IL-8 (Figure 3.a.) of about 27% were seen in the 3-in-1 heterocellular spheroid followed 21.43% in the Panc-1:J774.A1 spheroid. Comparatively low levels (20.8%) were seen in the Panc-1 homocellular spheroid followed by 18% in the Panc-1:NIH/3T3 spheroid. Numerous sources cite the role of TNF-a in the pancreatic tumors to be that of a double-edged sword, due to both, its pro and anti-tumorigenic effects. TNF-a levels are significantly elevated in pancreatic tumors and since they are primarily produced primarily by TAMs, the trend seen in the spheroid models was not surprising (41). Highest levels of TNF-α at 13% was observed in Panc-1:J774.A1 spheroid (Fig 3.b.). This was followed by comparatively low levels of 4%, 2.5% and 5% in the Panc-1 homocellular spheroid, Panc-1:NIH/3T3 spheroid and the 3-in-1 heterocellular spheroid respectively. Similarly, TGF-b has anti-tumorigenic effects at the early stages of tumor growth but at latter stages promotes tumor cell proliferation, desmoplasia and metastasis (42). Since TGF-b is produced by both tumor and stromal cells, the trend seen across spheroid models in our PCR experiment agreed with expected results (43). Highest levels of TGF-β (Figure 3.c.) of 56% were observed in the Panc-1:NIH/3T3 spheroid followed by 42% in the Panc-1 homocellular spheroid. It is interesting to note here that the Panc-1:NIH/3T3 spheroids were among the largest and densest, suggesting a morphological as well as a phenotypic relevance. Low levels of 26% and 18% were observed in the Panc-1:J774.A1 and 3-in-1 heterocellular spheroid, respectively. Although none of these results were statistically significant, the trends of their expression across different spheroid models are representative of the contribution of the different cell types.

**Figure 3.**
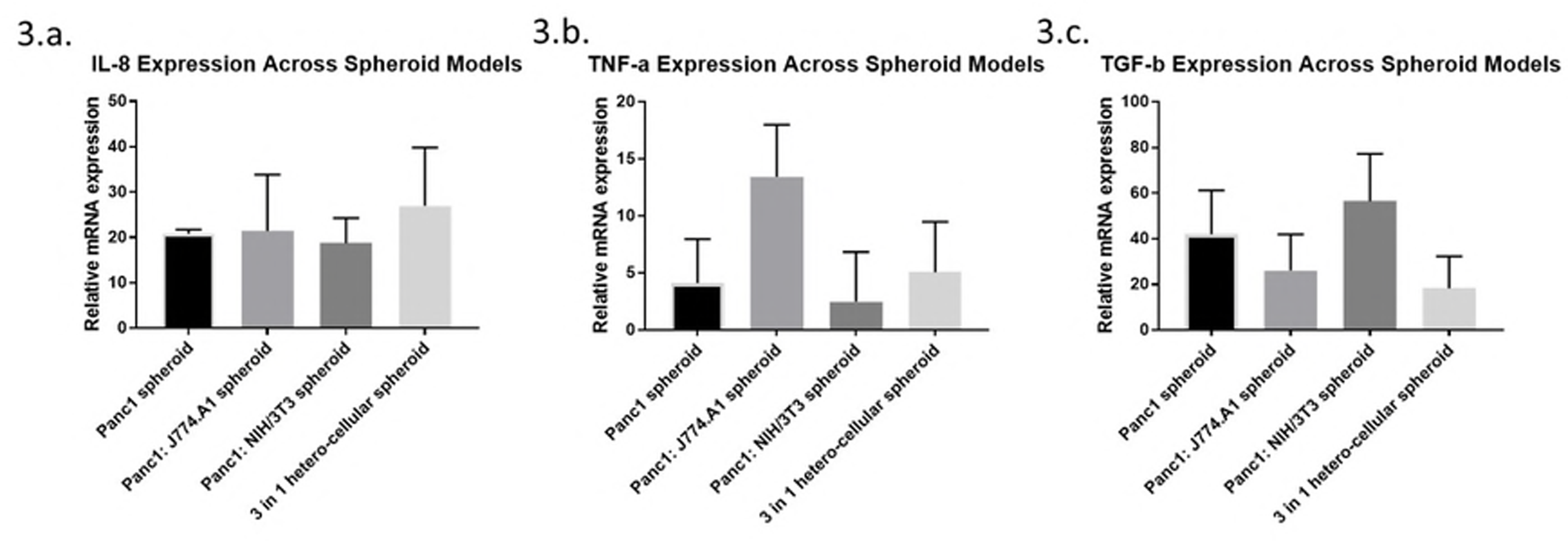
PCR evaluation of inflammatory markers in spheroid models. A. Statistical evaluation of expression of the IL-8 gene; B. Statistical evaluation of expression of the TNF-α gene with 1-tailed test across all spheroid models. C. Statistical evaluation of expression of the TGF-β gene. All expressions were normalized to the respective β-actin as housekeeping gene. The p values have been calculated to test differences compared to Panc-1 spheroids at significance limit of α = 0.05 using 1 way ANOVA.

TAMs and tumor cells under hypoxic conditions employ the glycolytic pathway and up-regulate HIF-1a (41). Analysis of qPCR results for HIF-1α revealed highest levels of expression in the Panc-1:J774.A1 spheroid at 1.5% compared to Panc-1 cells grown in normoxic conditions followed by that in the 3-in-1 heterocellular spheroids at 0.94% increase. Both results were statistically significant. p-Values were calculated using 1 way ANOVA and comparing values of spheroids with those in normoxic cells and tested at a significance level of α = 0.05. These results strongly correlate with the expected phenotype observed in pancreatic tumor microenvironments (41). The Panc-1:NIH/3T3 spheroid expressed HIF-1a at 0.43% and the Panc-1 homocellular spheroid at 0.27%. However the result for the Panc-1:NIH/3T3 spheroid and the Panc-1 homocellular spheroid did not reach statistically significance.

Due to increased hypoxia and glycolytic metabolism due to the Warburg effect, there is an associated low pH tumor microenvironment and an increase in HIF-2a expression (44). This increased expression is associated with dense stroma and so we hypothesized that the fibroblast containing spheroids may exhibit hypoxia like molecular feature. Indeed, qPCR analysis of HIF-2α level (Figure 4.b.) reveals highest levels of 147% in the Panc-1:NIH/3T3 spheroid followed by 15% in the 3-in-1 heterocellular spheroid (45). Both results were found to be statistically significant. Third highest HIF-2α levels were observed in the Panc-1:J774.A1 spheroid at 8.95%. Levels in normoxic and hypoxic monolayers were 6.27% and 5.15% and at 4.34% in the Panc-1 homocellular spheroids. These results assert that the multicellular spheroids are able to recapitulate the hypoxic microenvironment of a tumor, which is a hallmark property of cancer.

**Figure 4:**
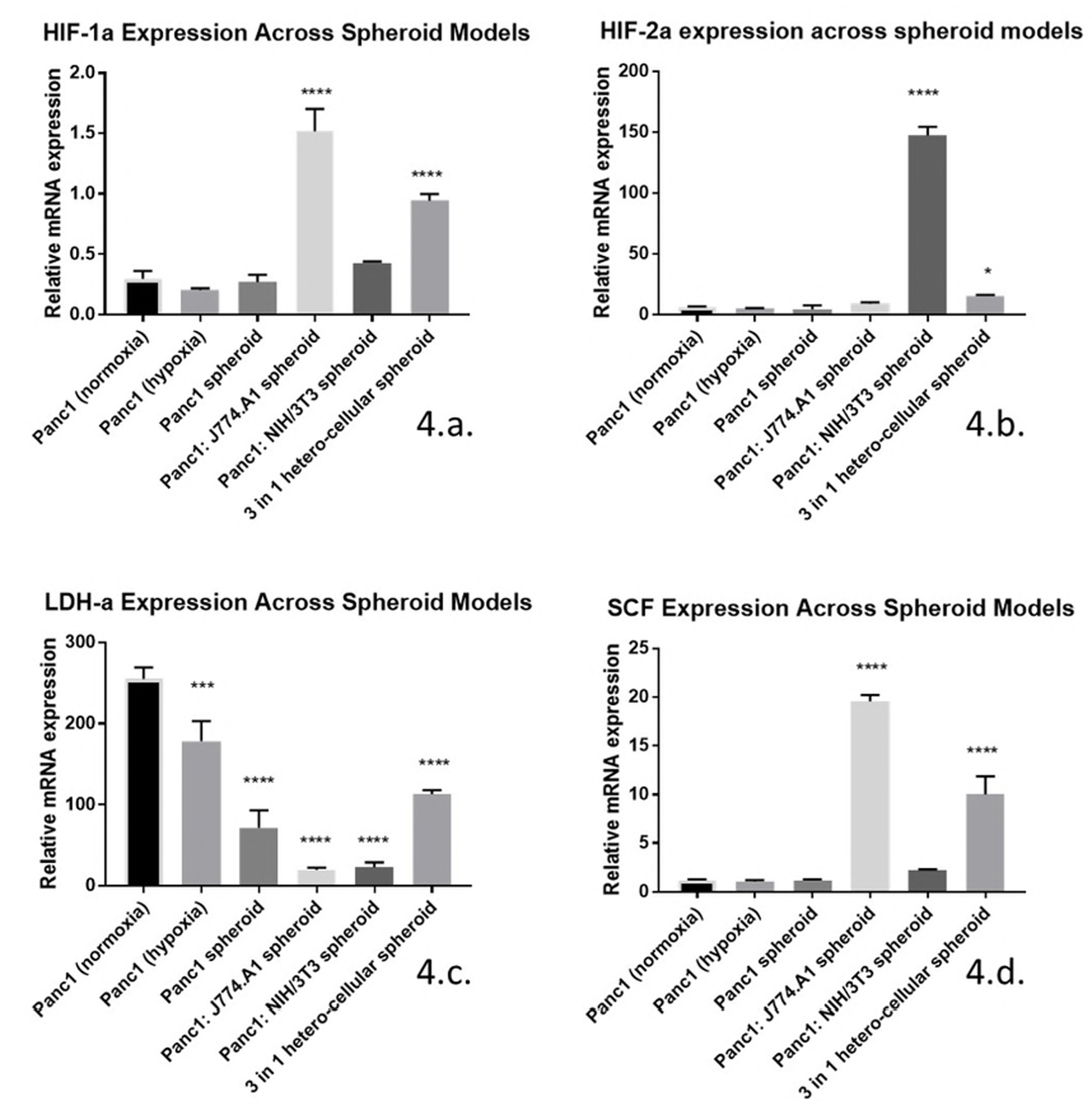
PCR evaluation of hypoxic and stem cell markers in spheroid models. A. Relative expression of HIF1α across different models. B. Relative expression of HIF-2α across different models. C. Relative expression of LDH-A across different models. D. Relative expression of SCF across different models. The p values have been calculated using one way ANOVA to test differences compared to Panc-1 cells grown in normoxic conditions at significance limit of α = 0.05

LDH-A levels were highest in the normoxic Panc-1 cells, as shown in Figure 4.c., at 255%, followed by 178% and 113% in hypoxic cells and the 3-in-1 heterocellular spheroid, respectively. Although we would have expected to see highest levels of LDH-A in hypoxic Panc-1 cells, hypoxic cells very quickly revert their phenotype to normoxic when exposed even briefly to oxygen. Since these cells were withdrawn from the hypoxia chamber before processing PCR samples, the brief exposure to oxygen may have obfuscated results seen in the monolayer samples (46). It is however interesting to note that the highest levels among spheroids were seen in the densest 3-in-1 heterocellular spheroid. The Panc-1 homocellular spheroid, Panc-1:NIH/3T3 spheroid and Panc-1:J774.A1 spheroid expression was 71%, 22% and 19% respectively. All results were found to be statistically significant. Binding of SCF to its ligand, c-kit, induces the up-regulation of HIF-1a through activation of PI3K/Akt and Ras/MEK/ERK pathways (47). Highest levels of SCF, shown in Figure 4.d., of 20% were seen in the Panc-1:J774.A1 spheroid, followed by 10% in the 3-in-1 heterocellular spheroid. These results were found to be statistically significant. Next highest levels were seen in the Panc-1:NIH/3T3 spheroid at 2.2%. Differences between SCF expression levels in the normoxic, hypoxic and homocellular spheroid were minimal and with no statistical significance. It is extremely interesting to note that correlation between HIF-1a and SCF expression across our models is suggestive of SCF induced HIF-1a upregulation. Immunostaining experiments for CD24 and SCF, both surface markers of cancer stem cells (CSCs), and subsequent imaging of fluorescence using the confocal microscope revealed presence of CSCs in Panc-1 homocellular, Panc-1:J774.A1 and the 3-in-1 heterocellular spheroid, shown in Figures 5 and 6. Detection of stem cell populations in these spheroid models through immunofluorescence along with the trends seen in inflammatory and hypoxia markers through qPCR further support our argument for using co-culture spheroid models as physiologically relevant testing platforms for anti-cancer therapies.

**Figure 5:**
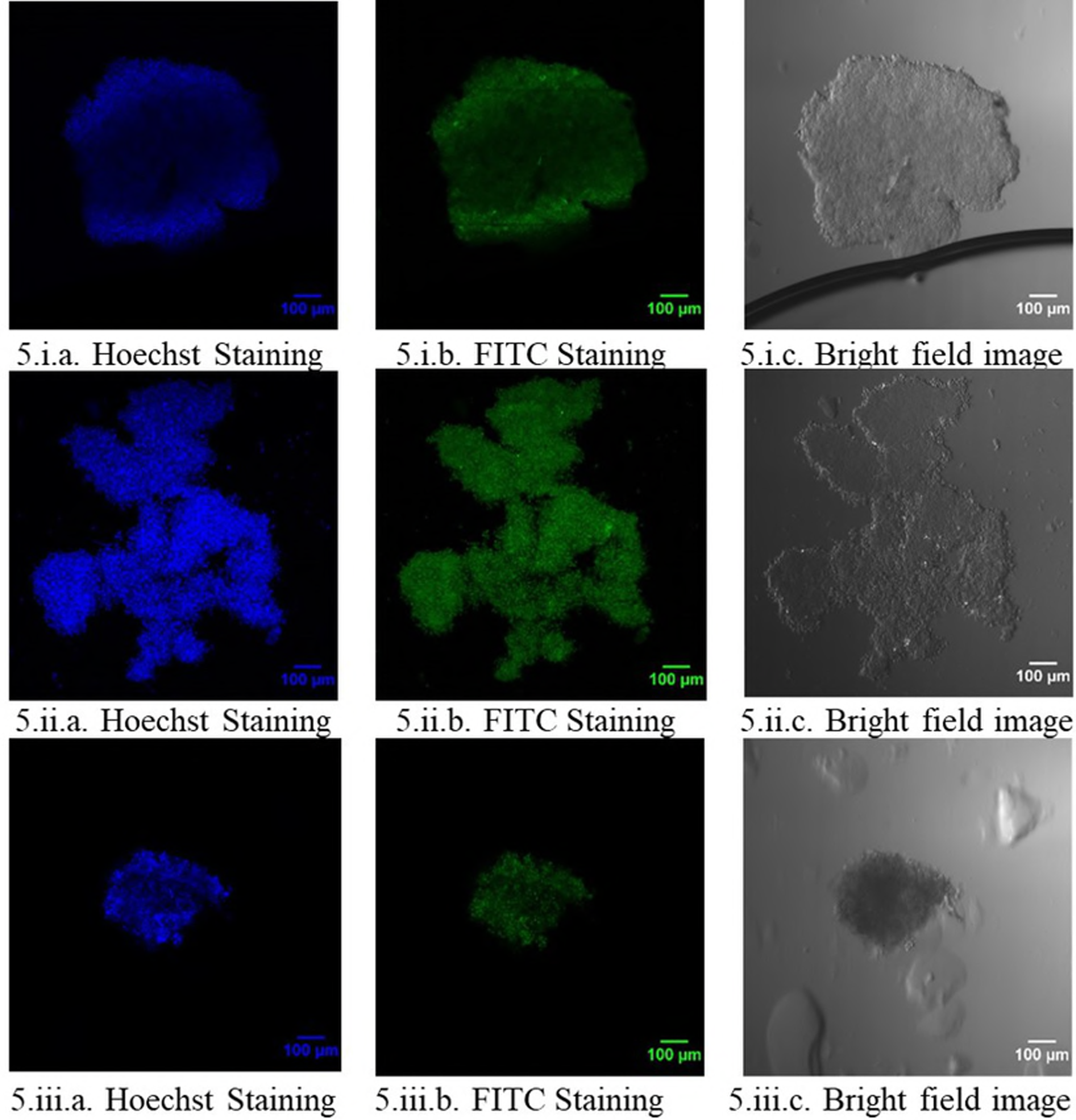
Immunofluorescence study of CD24 expression in spheroid models. i. CD24+ staining in Panc-1 Day 5 homocellular spheroids; ii. CD24+ staining in 1:1 Panc-1:J774.A1 Day 5 heterocellular spheroids; iii. CD24+ staining in 1:1:1 Panc-1:J774.A1:NIH3T3 Day 5 heterocellular spheroids

**Figure 6:**
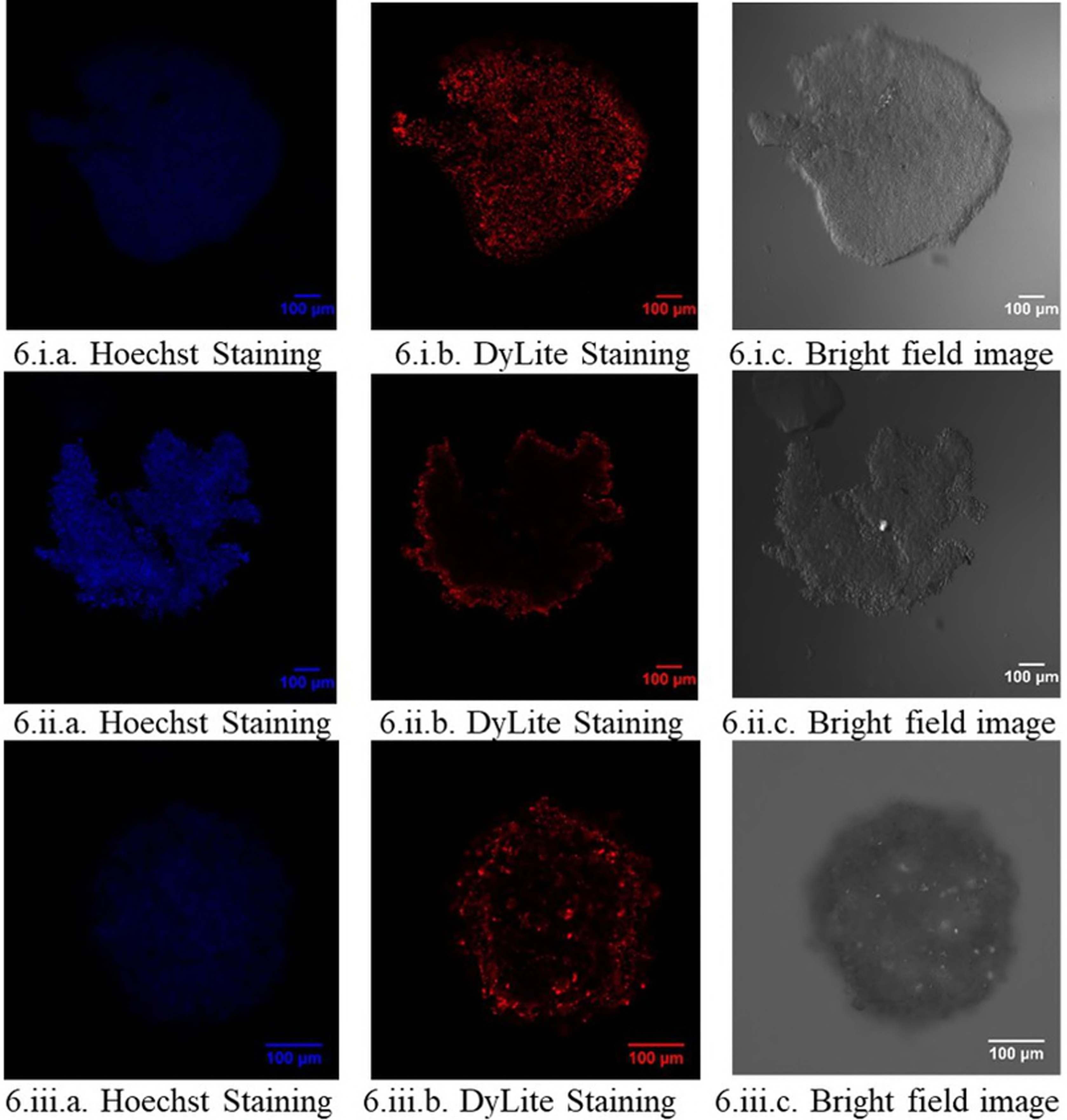
Immunofluorescence study of SCF expression in spheroid models. i. SCF staining in Panc-1 Day 5 homocellular spheroids; ii. SCF staining in 1:1 Panc-1:J774.A1 Day 5 heterocellular spheroids; iii. SCF staining in 1:1:1 Panc-1:J774.A1:NIH3T3 Day 5 heterocellular spheroids

## 4. Conclusions

The hanging drop method was used to successfully and reproducibly generate all four self-aggregating spheroid models, which were then characterized for their morphology and size. Pancreatic tumors are known to have a desmoplastic stroma, which not only provides structural support for the tumor as well as nutrients for the tumor cell proliferation, but also obstructs chemotherapeutics from reaching the tumor core. The random arrangement of cells in 3 in 1 heterocellular spheroids were indicative of proximity of each cell line with the other, and increased cell-cell interaction, as seen in the tumor microenvironment. Additionally, this model has the smallest diameter and depth, further supporting our argument for a stronger resemblance of pancreatic tumor cells in their physiological state.

Although analysis of the inflammatory markers IL-8, TNF-a and TGF-b through PCR did not reveal statistically significant differences across spheroid models, the trends in the expression suggests that the microenvironment of these spheroids is conducive to desmoplasia, cell proliferation, EMT and metastasis.

Hypoxia drives angiogenesis and EMT, contributes to MDR and enables cancer cells to switch their metabolism to the anaerobic pathway. Analysis of glycolytic and cancer stem cell markers through qPCR revealed much greater differences in expression levels in the 3D model compared to Panc-1 cells monolayer cultures. Highest levels of HIF1-a and HIF-2a seen in the co-culture spheroids indicate a stronger hypoxic component in these models, within which cancer stem cells can maintain their state of self-renewal, and prevent damage from ROS.

CSCs are pluripotent, immortal and can modulate their phenotype based on environmental cues. They are also resistant to the action of conventional chemotherapy and can self-renew. Positive staining of CD24 and SCF surface markers through immunofluorescence is indicative of the presence of CSC within these spheroid models. SCF levels being significantly higher in the co-culture models, as evidenced in the PCR study, offer further support to the hypothesis that cancer stem cell formation is largely dependent on microenvironmental cells and cross-talk.

This model has the potential to replicate the barrier to drug delivery seen in tumors. Additionally, co-culturing of different cell types that form spheroids have shown significant effects in altering tumor phenotypes and influencing CSC population growth, thereby re-enforcing the need to develop robust and clinically relevant *in vitro* models for testing chemotherapeutic agents.

## Acknowledgements

Financial support was provided by the National Cancer Institute of the National Institutes of Health through grant R21-CA213114.

